# Context of Psychotropic Drug Delivery Modulates its Neurobehavioral Effects: the Case of Methylphenidate

**DOI:** 10.1101/212480

**Authors:** Roy Sar-El, Gal Raz, Nitzan Lubianiker, Haggai Sharon, Talma Hendler

**Author notes:** R.S-E. and G.R. contributed equally to this study. **Corresponding author:** Nitzan Lubianiker.

## Abstract

Pharmacotherapy is substantially hindered by poor drug targeting, resulting in low specificity and efficacy. Here, we tested a novel, non-invasive targeting approach (termed functional-pharmacology), which couples drug administration with a task that activates the drug’s sites-of-action in the brain, thus possibly improving absorption and efficacy. Methylphenidate (MPH) or Placebo were administered to healthy subjects, which then performed a cognitive induction or a control task. N-Back fMRI before and after drug-task coupling measured therapeutic effects. Only following MPH, subjects that performed better in the cognitive induction task showed greater improvements in N-back performance. Moreover, only under MPH-Cognitive induction condition, there existed a significant correlation between improved recruitment of N-Back rDLPFC activation, and a concurrent improvement in task performance. Importantly, mediation analysis suggested a causal role of rDLPFC activation in these coupling effects. Our results support the functional-pharmacology concept feasibility and efficacy, hence opening a new horizon for patient-tailored, context-driven drug therapy.

## Introduction

Efficacy of psychopharmacological treatment is often considerably restricted by the fact that drugs reach both pathologically relevant as well as irrelevant brain areas in a nonselective manner, thus causing desired but also unwanted effects. A bedside method of enhancing specificity of drug delivery would have important clinical implications. First and foremost, it may allow a reduction in the dosage required for satisfactory therapeutic effects in the target area (i.e. greater efficacy), resulting in reduced abuse and improved side effect profile. In addition, and no less important, it may result in better adherence and reduced economic burden.

In an effort to tackle the need for better drug targeting, researchers have thus far employed a range of different technological means such as nanocarrier administration [1][2], laser stimulation [3], ultrasound [4], and polymer implantation[5]. However, these are all expensive, complicated and often invasive procedures. An alternative approach may be based on the idea that contextual physiological and psycho-physiological factors significantly interact with drug effect profiles. This rationale is analogous to the consideration of the gastrointestinal "context" for better systemic drug absorption, e.g. taking a drug before or after a meal, in the morning or evening.

In this study, we demonstrate a novel approach for non-invasive drug targeting, based on the assumption that therapeutic drug effects in the brain may be enhanced by inducing a specific advantageous physiological state in the drug-target during its absorption and distribution. In other words, coupling drug administration with functionally relevant local neural activation (termed hereby; *Functional Pharmacology*). Such selective activation of a targeted brain region or circuit would potentially result in desired modulation of key factors that determine drug delivery effects such as pharmacokinetics and pharmacodynamics.

For example, the cerebral blood flow (CBF) is a key factor affecting the movement and diffusion of drugs in the brain (i.e. pharmacokinetics), influencing the amount of the drug which reaches the Blood Brain Barrier (BBB) and ultimately arrives to its target sites [6] [7]. Crucially, focal increases in neural activity in the brain are accompanied by increased regional blood supply (for a review, see [8]) with an estimated magnitude of 47% - 60%, [9], [10] in different sensory processing areas. This effect relies on the neurovascular coupling – a process in which neurons, astrocytes, and vascular cells interact to create local hemodynamic changes. Recent work has shown that neural activity can also modify neurotransmitter expression in the brain leading to re-specification of receptors (i.e. affecting drug Pharmacodynamics) [11]. This effect is known to be context dependent [12].

To demonstrate our new approach, we selected Methylphenidate (MPH), an inhibitor of monoaminergic reuptake, commonly used to treat attention deficit hyperactivity disorder (ADHD). Importantly for this approach, MPH is flow dependent, a characteristic that is measured by the difference between the arterial and venous drug concentration (which is high in MPH). This means that it is extracted from the circulation rapidly, accumulating in the BBB and eventually increases the amount that enters the brain [7], [13].

While the specific mechanism by which MPH enhances cognitive performance has yet to be fully understood, accumulating evidence points to the importance of its influence on the interplay between the mesocortical and the mesolimbic pathways in the brain (for a review, see[14]). MPH allows demonstration of differential targeting of the dopaminergic system by functional local activation. Dopamine in the mesocortical pathway (from the Ventral Tegmental Area (VTA) to the Prefrontal cortex) plays an important role in cognitive and executive functions including working memory and underlies MPH's effect on attention (for a review, see [15]. However, Dopamine in the mesolimbic pathway (ventral tegmental area (VTA) projections to the ventral striatum) has a crucial role in the reward system, and is responsible to MPH's side effects on emotional modulation [14].

It is well known that behavioral context can facilitate changes in the local cellular reactivity [12]. We therefore set out to examine whether the coupling of MPH administration with a cognitive task (Cognitive-MPH) would result in increased drug effect on cognitive performance that is known to involve prefrontal cortex activation. To test the specificity of the effect of Cognitive-MPH, we added a control task induction condition in which MPH administration was coupled with a task known to activate mesolimbic areas (Control-MPH) [16]. To control for overall task effects (beyond the drug effect) we added a placebo drug condition for each task (Cognitive-Placebo, Control-Placebo).

Before and after all four conditions, a commonly used executive function N-Back task was performed during fMRI. Since MPH treatment in ADHD is related to increased activation in the prefrontal cortex [17], and since the right DLPFC (rDLPFC) was specifically shown to be related to cognitive load in executive function [18], [19] such as in the N-Back task [20], it was a-priori defined as a region of interest (ROI) for our fMRI indication for neural change along with the cognitive performance change. Accordingly, we expected to see the greatest change in N-Back task activity in the ROI following Cognitive-MPH, in comparison to the other conditions

We specifically hypothesized that: **1.** Selectively coupling MPH administration with a cognitive task known to best activate the rDLPFC, would result in greatest improvement in executive function performance, as measured by the N-back task during scanning, compared with the other conditions (i.e. Placebo-Cognitive, MPH-Control, Placebo-Control); **2.** The improved performance during scanning would correlate with the change in task activity in rDLPFC following drug administration, only in the MPH-Cognitive condition; **3.** The increased activity in the rDLPFC following drug administration in the MPH-Cognitive condition would correlate with the performance in the coupled cognitive induction task.

In order to test these hypotheses, participants underwent four experimental sessions in a double-blind, counterbalanced, within-subject factorial design, with sessions interspersed by at least a week. In each session, participants received either 30 mg MPH (P.O) or an identical looking starch pill as a placebo. Previous studies which used oral MPH have shown that 20-40 mg corresponds to ~50% dopamine receptor occupancy [13], [21], [22] a suitable dose to avoid ceiling effect. Drug administration (MPH or Placebo) was immediately followed by either a cognitive or a control task, resulting in four conditions: MPH-Cognitive, MPH-Control, Placebo-Cognitive, and Placebo-Control (See Figure 1). Both tasks were applied in a quiet examination room and lasted each approximately 45 minutes. All sessions were conducted at approximately the same time in the morning. Participants were instructed to refrain from consuming caffeine and alcohol for 24 hours prior to each session and this was verified verbally at the beginning of each session.

**Figure 1:**
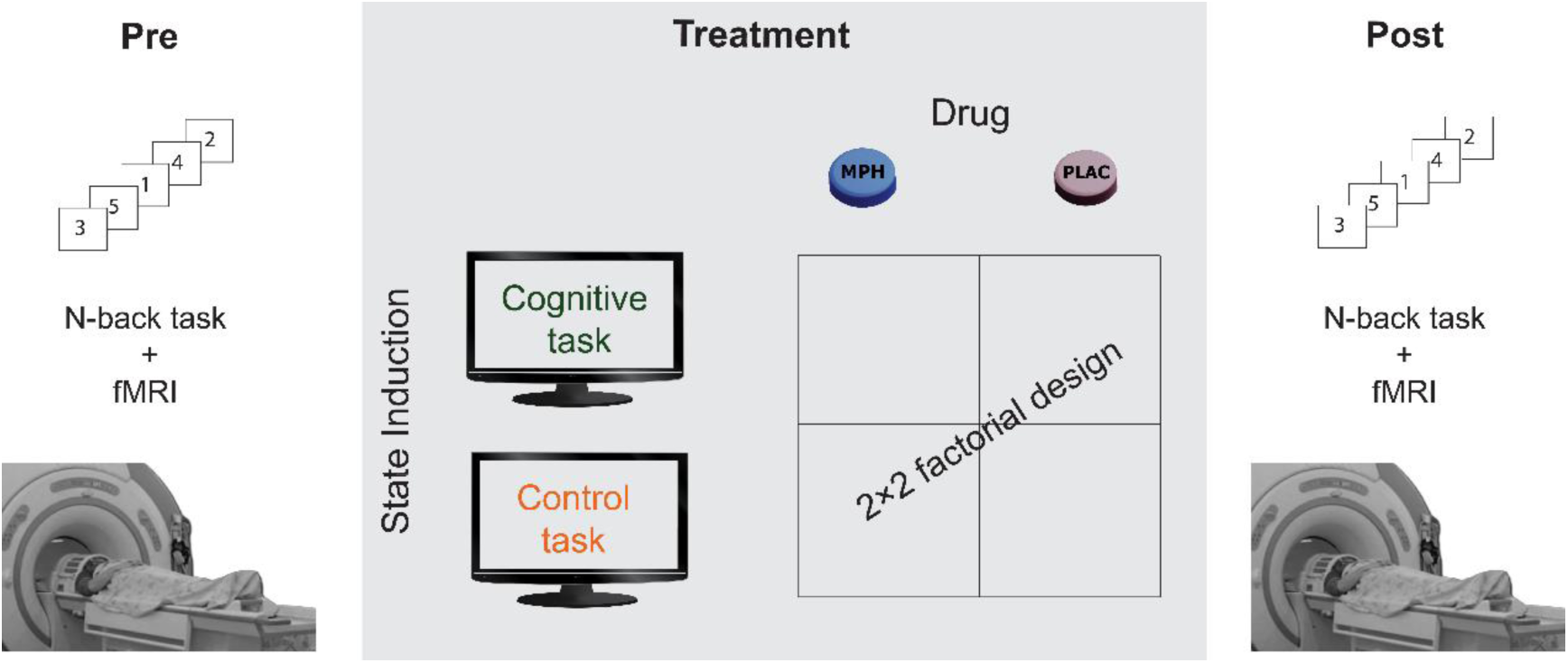
A schematic description of the experimental design. Participants underwent four functional pharmacology counterbalanced sessions in a within-subject 2×2 factorial design, resulting in four coupling conditions: Cognitive-MPH, Cognitive-Placebo, Control-MPH and Control-Placebo. fMRI scanning during the N-Back task was performed before the drug-task coupling procedure and immediately afterwards.

For the cognitive state induction we used a validated battery of 7 computerized cognitive tasks (NeuroTrax Corp., Bellaire, TX), known to probe executive functions, such as GoNoGo and Stroop interference tasks [23]. Test scores for accuracy and response time (RT) were normalized to a standard scale with a mean of 100 and standard deviation of 15, based on normative data from a large cohort (n=1569) of cognitively healthy individuals provided by the NeuroTrax manual. A global cognitive score was computed as the average of all test scores for a single administration of the cognitive battery. In addition, an "attention index" was computed based on the RT in attention tasks sensitive to ADHD: GoNoGo, staged information processing and stroop interference[24]. As the control state induction, we employed two paradigms known to probe reward processing and mesolimbic activation. One is a 25 minute competitive computer game developed in our lab as a paradigm to assess goal directed behavior with regard to reward and punishment [16], and the other is a 20 minute passive listening excerpt using emotional music extracted from the Pachelbel's Canon, previously used in our lab to induce positive emotions [25].

The N-Back task was performed during fMRI scanning before and 60-90 minutes following the drug-task coupling procedure on each experimental day. The time interval between drug administration and scanning corresponds to the expected time-to-peak brain concentration of MPH after a single oral administration [26], [27].

The relation between brain activity and behavioral performance during high cognitive load was examined for each session by correlating the change in RT (msec) with the corresponding change in the rDLPFC activity (beta values) during the 3-Back condition. The relation between the cognitive induction and the functional-pharmacology behavioral outcome was tested only for Cognitive-MPH/Placebo sessions, by correlating the change in 3-back RT (pre-post) during scanning with the performance on the cognitive induction task that was coupled with administration of the drug (MPH or Placebo). Various correlations were compared using the Steiger's Z-test for dependent correlations.

To gain further insight regarding the mechanism of functional-pharmacological coupling, we applied a mediation analysis following a standard three variable path model according to the INDIRECT procedure of SPSS [28]. The mediation analysis was performed on three variables from the MPH-Cognitive condition: the change in 3-back RT (pre-post) during scanning; the corresponding change in rDLPFC activity (beta values) during the 3-back period; and the "attention index" obtained from the cognitive induction task that was coupled with MPH. The indirect effect was significant if its 95% bootstrap confidence intervals from 10000 iterations did not include zero at p=0.05.

## Results

The validity of *functional-pharmacology* procedure was evaluated by the performance on each task coupled with either MPH or placebo. Cognitive performance on executive functions tasks was measured according to the NeuroTrax attention index, which did not differ between MPH and Placebo (97.216 and 100.441, respectively, t(12)=-1.04, p=0.32), suggesting that the cognitive induction was similarly effective in both drug conditions. In the control task, the motivation index was calculated as the improvement in game score from session 1 to session 4. Game scores were higher in the MPH condition than in the placebo condition (402.167, 282.833, respectively, t(12)=3.0828, p=0.01), suggesting an advantage for MPH in this task.

The effect of *functional-pharmacology* on drug targeting was evaluated by assessing the N-back performance (RT and accuracy) and rDLPFC activity. We first examined whether there existed any differences between conditions in performance prior to all drug-task couplings. There were no differences in the 3-back RT (F(3,36)=0.331, p=0.803) and in the 2-back response time (F(3,36)=0.207, p=0.891) between the four conditions. Then, we validated the cognitive-load effect of the N-Back task across drug conditions and time points in behavioral and brain measures. There was an expected behavioral effect of cognitive-load, both on RT and accuracy measures across drug conditions and time points. The RT for 0-2-3-back was 437.35±16.67, 553.79±43.13 and 645.49±61.19 msec, respectively (repeated measures ANOVA, F(2,24)=7.456, p<0.01, 0 vs 2 back p<0.05, 0 vs 3 back p<0.01, 2 vs 3 back p<0.05, bonferroni-corrected). The mean accuracy for the 0-2-3-back was 96.47±1.5, 86.00±3.21 and 68.57±5.36 msec, respectively (repeated measures ANOVA F(2,24)=16.832, p<0.001; 0 vs 2 back p<0.01, 0 vs 3 back p<0.001, 2 vs 3 back p<0.001, bonferroni-corrected). These behavioral results confirmed that the N-back task indeed manipulated working memory with differential cognitive load effect.

For a similar cognitive load effect in brain activity, we first obtained whole-brain activation maps, across drug conditions and time points, for the 3-back versus 0-back. As expected, there was greater activity for 3-back than 0-back (p<0.01, FDR-corrected), encompassing typical executive function activity in fronto-parietal areas including the right DLPFC (Figure 2a). ROI analysis for the rDLPFC further showed this effect (Repeated measure ANOVA F(2,24)=74.11, p<0.001), with 0-back showing lower activity than 2-and 3-back, for each time point (p<0.001 for both, bonferroni-corrected) (Figure 2b).

**Figure 2:**
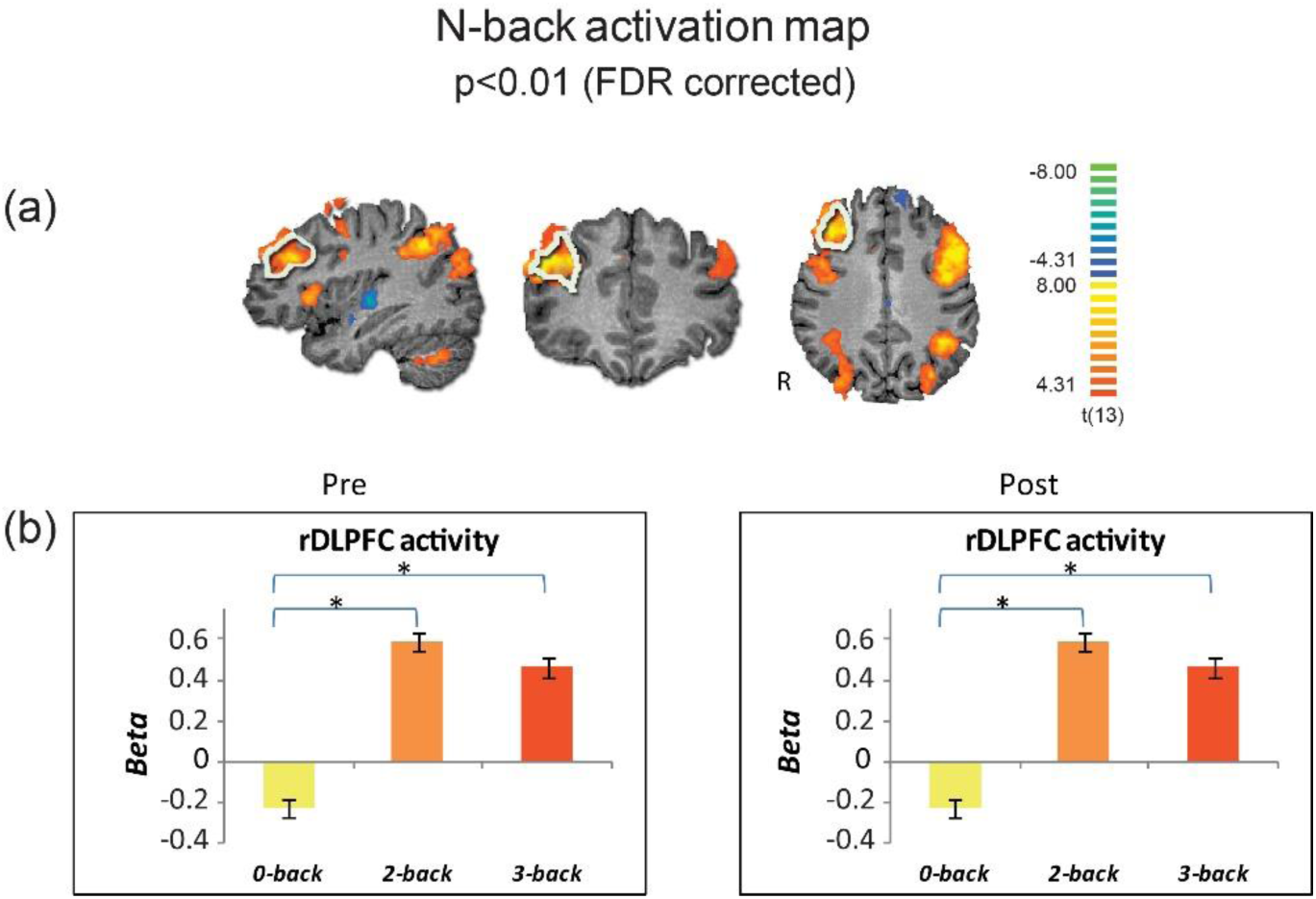
Cognitive load effect on brain activity. (a) Group brain activation maps (n=13) for 3-back vs 0-back are shown in sagittal coronal and axial views (random effects, p<0.01, FDR corrected). The a-priori selected ROI in the rDLPFC is marked by white boundaries. (b) Activity in the rDLPFC shows similar cognitive load-effect for pre-and post drug administration across induction conditions. Error bars stand for Standard Error of the Mean. *=p<0.001

To assess drug specific effect on the *functional-pharmacology* procedure we first looked for changes in the N-back task performance (RT and accuracy) and rDLPFC activity (beta values), using a 2-way repeated-measures ANOVA with drug (MPH/Placebo) and induction task (cognitive/control) as factors during the 3-back load condition. Neither main effect for induction task or drug, nor interaction between factors were found for behavioral (drug main effect: F(1,12)=0.234, p=0.637, induction task main effect: F(1,12)=0.004, p=0.953, drug X task interaction effect: F(1,12)=2.637, p=0.130) or neural measures (drug main effect: F(1,12)=1.5, p=0.244, induction task main effect: F(1,12)= 0.018, p=0.895, drug X task interaction effect: F(1,12)=0.6430, p=0.443).

We then examined individual variability for drug effect by correlating between RT and rDLPFC activity during the 3-back load condition, per *functional-pharmacology* coupling condition. Figure 3 shows that only during the MPH-Cognitive condition there was a significant correlation between change in rDLPFC activity (Post>Pre drug) and RT (Post<Pre drug), indicating that greater increase in rDLPFC activity corresponded with greater reduction in RT following MPH (R=-0.75, p=0.003; left upper panel). Steiger's Z-test indicated a significant difference between the correlations and a unique association only for the cognitive-MPH condition (Steiger’s Z=-2.934, p<0.05, bonferroni-corrected for multiple comparisons).

**Figure 3:**
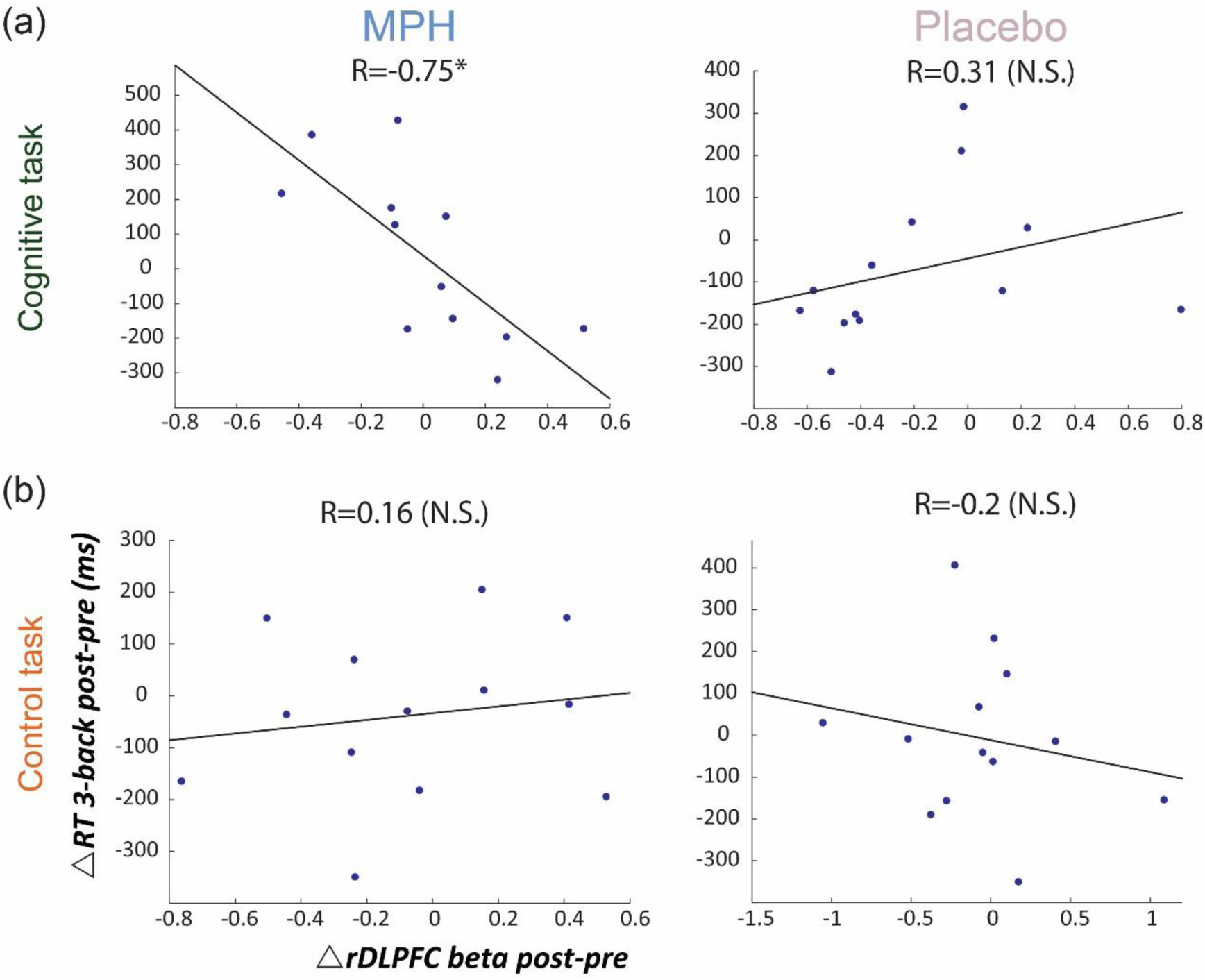
Brain-behavior correlations for the 3-back load, with respect to drug administration and coupling condition. Relationship between change in RT (post vs pre) and rDLPFC activity (post vs pre) during the 3-back, following each coupling intervention (MPH-Cognitive, Placebo-Cognitive, MPH-Control and Placebo-Control). Only the MPH during cognitive task revealed a significant correlation (r=-0.75, P<0.01; Steiger’s Z=-2.934, p<0.05, bonferroni-corrected for multiple comparisons).

To further validate the coupling effect of *functional-pharmacology*, we examined whether the behavioral improvement following drug-task coupling corresponds to subjects’ performance during the cognitive induction task, and whether this is evident specifically in the MPH condition. This follows the assumption that given the presence of MPH in the brain, better performance during cognitive induction would recruit cognitively relevant brain regions and result in greater improvement in the N-Back working memory task. In line with this, Figure 4a shows that the attention-index derived from the Neurotrax battery was higher among individuals who showed higher improvement in RT on 3-Back following MPH (r=-0.76, p<0.01), but not placebo (r=0.03, p=0.97). Steiger’s Z-test for dependent correlations indicated a significantly stronger correlation for the MPH-Cognitive state coupling relative to the Placebo-cognitive coupling (Z=-3.34, p<0.001). Similar results were found for NeuroTrax global score (MPH, r=-0.69, p<0.01, Placebo; r=0.32, p=0.24). To further demonstrate this coupling effect at the individual level, we classified subjects into groups of high and low attention index scores (Figure 4b) and present their 3-back RT change during scanning (i.e. indicated by colored lines connecting pre and post RT measures). As expected, following the MPH condition, individuals from the high scoring group (n=7) showed greater improvement in the 3-back RT, as indicated by the existence of more red colored lines. On the contrary, those from the low scoring group (n=6), in fact worsened following drug administration (RT was slower) as indicated by the existence of more purple connecting lines (Mean Diff=-111.15±109.08 and Mean Diff=169.41±47.64, respectively; t(11)=2.487, p<0.05). Importantly, this difference was not evident during the placebo condition (not shown).

**Figure 4:**
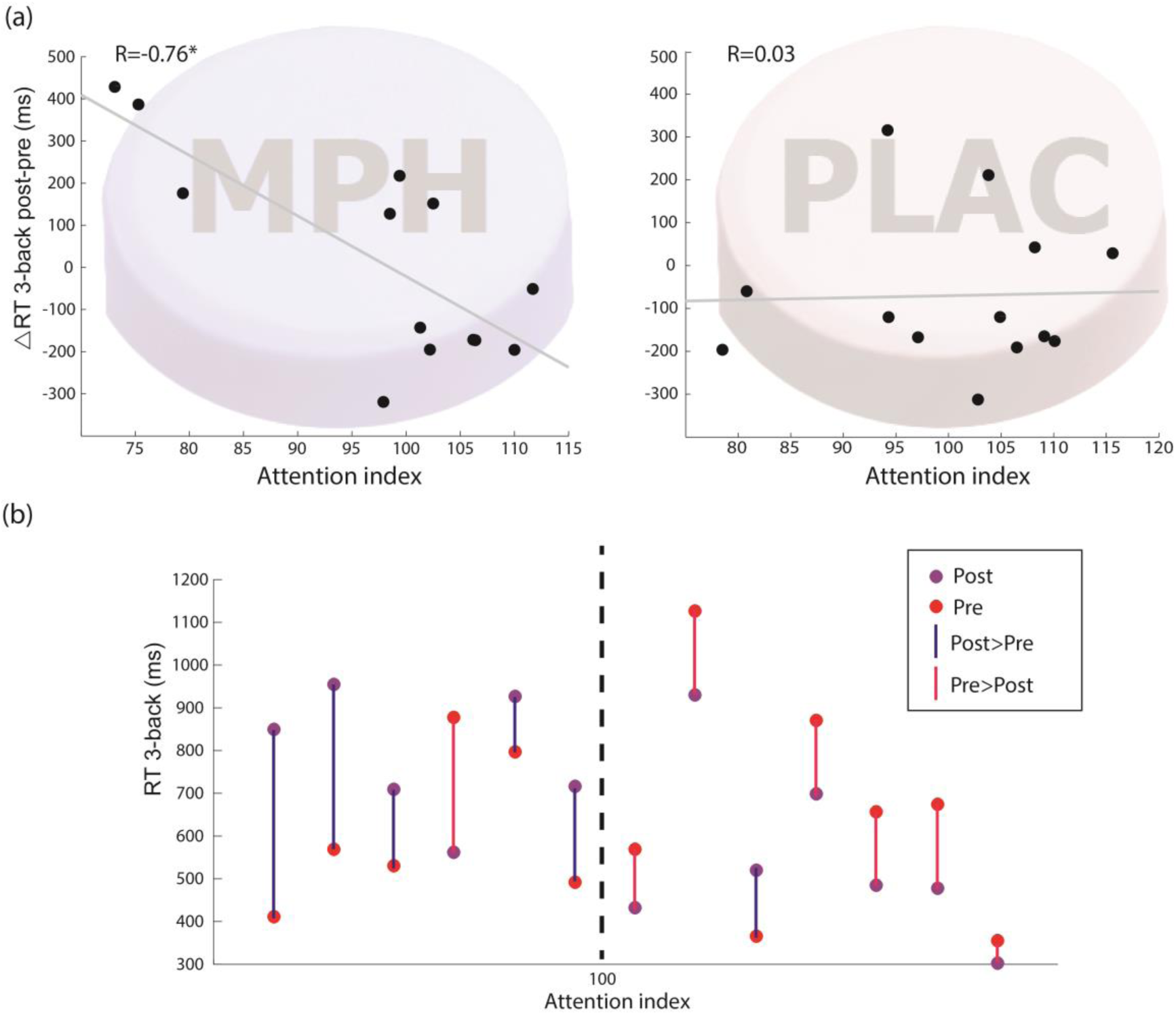
Relationship between performance during the cognitive induction task and improvement in N-back performance during brain scanning. (a) Correlation between attention index during the induction phase and 3-back RT difference (post vs pre drug-task coupling). Performance on cognitive induction task coupled with MPH, but not with placebo, showed a significant correlation with 3-back RT change during scanning (R=-0.76; p<0.01). (b) Individual measures of the correlation between performance during cognitive induction and the change in 3-back RT during scanning for the MPH-Cognitive condition. Individuals are clustered into high and low induction performance according to the attention index (score above or below the population average as indicated by a dashed line, NeuroTrax Corp., Bellaire, TX). The RT measures for pre and post drug-task coupling are marked by red and purple dots, respectively. The difference in direction is marked by colored lines connecting the dots (purple for post>pre, and red for pre>post). Most individuals (6/7) in the high attention index during the cognitive induction (right to the dashed line), showed faster RT following MPH administration (red line connects the dots).

Finally, to assess causal relations in the observed association between change in the rDLPFC activity and RT in 3-back condition following MPH, we performed a mediation analysis [28] with the attention index of the cognitive induction as a mediator. As expected, we found a significant indirect path from rDLPFC activity and RT improvement through the performance during the cognitive induction coupled with MPH administration (indirect effect= -357.6, SE= 198.67, CI (95%) = -766.1 to -10.7) (Figure 5). This effect was not found for the placebo condition (indirect effect= -24.26, SE= 84.39, CI(95%)=-450.78 to 73.51), Suggesting attention scores during cognitive induction coupled with MPH mediated the association between rDLPFC beta difference and 3-back RT difference.

**Figure 5:**
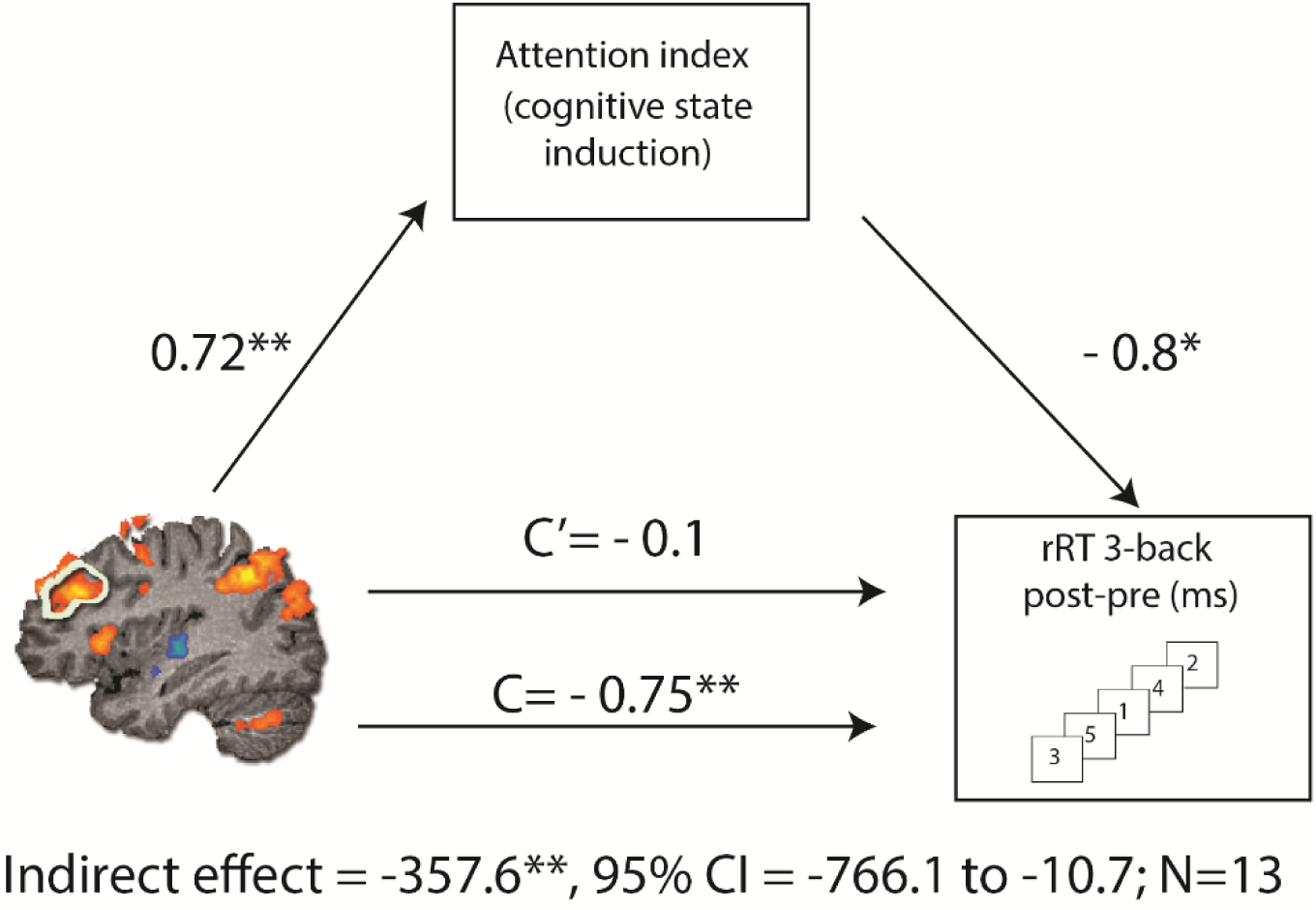
Mediation analysis describing possible causal mechanisms by examining variables that partially or fully account for the relationship between two variables. C represents the direct effect of rDLPFC beta difference on change in RT for the 3-back condition, while C' represents the indirect effect, when including the neurotrax attention index in the model. Including the attention index in the model results in a non-significant effect (C'), consistent with the mediation hypothesis. The indirect effect was tested using a bootstrap approach with 10000 samples. The model is considered significant since its 95% bootstrap CI from 10000 iterations did not include zero at p=0.05.

## Discussion

The current study introduces a novel concept for improved drug targeting in neuropsychiatry, namely the coupling of drug administration with a behavioral induction of a specific, functionally relevant, localized neural activity for the optimization of pharmacotherapy efficacy. As a proof-of-concept experiment in healthy individuals, we coupled the administration of MPH with a demanding cognitive task and tested drug targeting effects with executive function task performance and its related rDLPFC fMRI BOLD activity.

Taken together, our findings suggest that the coupling of MPH administration specifically with a cognitive induction task may improve performance (i.e. drug effect) in a subsequent working memory challenge (Figure 3). Moreover, this beneficial effect does not result from a passive and uniform process inherent to performing the actual task, but rather depends on the individual performance during the cognitive induction condition (as indicated by the attention index) (Figure 4). Our findings also advocate that the drug-functional task coupling promote a lasting facilitation of rDLPFC recruitment and a corresponding behavioral improvement in the context of high cognitive load (Figure 5). This mediation supports our assumption that the actual functionally relevant coupling may improve drug targeting.

Our findings are in line with the notion that neural activity may modulate the ever-changing state-dependent neuronal environment [11], [12], [29] and thus influence the effectiveness of pharmacotherapy at any given time point depending on the current functional activity and connectivity of its relevant target circuit.

Taking the case of dopamine and MPH as an example, a multitude of animal studies [30]–[34] has repeatedly demonstrated that the behavioral effects of dopaminergic psychostimulants may vary dramatically, depending on the environmental context in which they are administered. For example, in humans it was shown that dopaminergic effects of cocaine were amplified under certain conditions, such as performing a salient, novel and unpredictable task [35]. Similarly, a previous study [36] showed that MPH's effect on hyperactivity and inappropriate behavior was increased when given in the context of learning compared to playing.

The current study adds to this existing literature by systematically manipulating the coupling of either drug or placebo with a relevant/irrelevant task, hence causally modulating behavior while examining the neural correlates of such context-dependency. A behavioral change that is contingent specifically on the combination of cognitive task and MPH administration, along with a domain-relevant change in brain activation (increased rDLPFC activation during high cognitive load), suggest that the functional-behavioral coupling harnesses the therapeutic effect of the drug in the specific designated neural circuitry, rather than an overall increase in drug delivery through a compensatory or collateral mechanism. In other words, the brain-behavior correlation reported here provides evidence that relevant context manipulation may enhance drug therapeutic effects through beneficial relevant neural targeting. Future studies using PET and specific drug related ligands should further examine the relation of this effect to local receptor activity.

A bedside method of enhancing targeted drug delivery and efficacy would have important clinical implications. First and foremost, it may contribute to enhancing the desired effects of the medication by functionally targeting drugs to their sites of action. Additionally, it may enable clinicians to reduce the systemic dosage needed to attain the same neural effects. If this proves to be the case, there follow various crucial benefits: first, reduced doses may mean reduced costs of treatment; second, if doses can be reduced in terms of frequency of administration, it may lead to more convenient administration regimes, thus promoting better treatment adherence; third and most importantly, reduced doses may reduce side effects of many neuropsychiatric drugs. Another significant advantage in the case of centrally acting drugs is the potential reduction in abuse rates. In the case of chronic opioid treatment, for instance, there exists a clear association between daily opioid doses and mortality as well as addiction; the challenge is therefore to find mitigating approaches that will enable reduced doses of opioids given to chronic pain patients but with equal effect [37]. To this end, the functional pharmacological coupling, which enables to reduce doses while maintaining satisfactory therapeutic effects, can provide an optimal solution.

The multi-systemic and widespread effects of MPH in various brain loci and its complex neuropsychiatric effects (for examples regarding dopaminergic widespread effects and its connection to side effects, see [38]) provide an ideal example as to the need of better functional-anatomical selective targeting of psychopharmacological drugs. Although MPH is considered a relatively safe drug, its use nevertheless leads to various adverse effects such as sleep difficulties, decreased appetite, nervousness, increased heart rate and blood pressure [39], all of which emphasize the need for a more focused and individualized delivery. Furthermore, some severe adverse effects such as cardiac events have been associated with MPH use, although these are rare ([40], [41], There is a considerable MPH discontinuation rate, which is associated with both poor response and adverse effects [42]. Additionally, it also presents a potential for abuse and medical or recreational use, particularly in childhood and adolescence, thus enhancing abuse potential also for other recreational substances, such as nicotine and cocaine [43], [44]. Our results support the notion that a better targeted administration may enhance drug effects while reducing its side effects, either as a consequence of using lower doses, or by inducing a physiological state which diverts the active ingredient from side-effects related brain regions.

It is well accepted that neuronal engagement, by increasing metabolic demands, enhances local blood flow in the relevant brain region [45]. This provides an elegant and rather intuitive rationale for functional-pharmacological coupling as a means to increase local drug delivery. The scale of the change in local blood flow is therefore of critical importance. It is widely accepted that the total CBF may be significantly modulated as a measure of brain activation [46]. Local neurovascular coupling may contribute a further significant and sustained increase in local blood flow over peak drug delivery time and may prove to be substantial. Accordingly, direct measures of CBF during sensory stimulation have documented up to 100% increases in local CBF [47], and therefore suggest a major impact of behavioral modification of neuronal activity on regional blood supply. This dramatic change in delivery would of course be especially relevant for drugs with a high partition-coefficient index, an indicator of BBB permeability and as a consequence, of flow dependency [7]. In addition, inducing a more demanding metabolic state, i.e. activating neurons and supporting cells, may lead to a multitude of cellular and membrane synthetic and conformational changes, which might be beneficial in the context of specific drugs. This principle can be demonstrated, for example, by the pharmacologic blocking of voltage-gated sodium channels with anticonvulsant medications, which exert their effects on neurons in a state-dependent block [48].

Our study presents several limitations, which should be addressed in future studies. First, this was a proof-of-concept study with healthy participants. In the case of cognitive effects of MPH on attention and cognition, this results in a ceiling effect. A clinical study with ADHD patients would be expected to provide additional information and support for this proposed approach. Second, several relevant domains other than executive function/working memory may be probed in the context of the specific neuropsychiatric profile of MPH. Further research examining the effects of functional-pharmacological coupling on other cognitive functions such as response inhibition and sustained attention may prove revealing [49]. Lastly, this study explored the immediate effect of a single administration of MPH. A longer-term administration scheme and follow up may provide additional vital insights.

## Conclusions

The notion of functional-pharmacological-coupling opens a new horizon for medical applications considering the rapidly growing neuroscientific literature on specific stimuli that activate certain brain regions on the one hand and on disease relevant neural circuitry on the other hand. In this respect, it binds together the fields of experimental cognitive neuroscience and clinical implication of psychopharmacology. It presents the opportunity for a multitude of clinical benefits and opens the door to a vast array of further investigations into specific links between functional and chemical maps in the brain. Moreover, it closely relates to the ideal of personalized patient-tailored medicine, as functional-pharmacological coupling protocols may be individualized and optimized to fit a specific subject's neural pattern of activation or physiological state to provide better mechanism-based neuropsychiatric pharmacotherapy.

## Methods

### Participants

17 healthy participants were recruited to the study via advertisement in social media. Two participants were excluded due to technical problems. Two participants completed only two out of the four sessions due to personal reasons. Thirteen participants therefore completed the study (Male=10, age: 25.83 ± 5.46; for all statistical analyses n=13). Participants reported no current use of psychoactive medications or illicit drugs, and had no family history of major neurological or psychiatric disorders. All participants provided written informed consent approved by the Tel Aviv Sourasky Medical Center's Ethics Committee. Participants were screened by the Adult ADHD Self-Report Scale (ASRS); six questions regarding ADHD symptoms, providing a summary score of 0-24, where 14-24 indicates ADHD [50], [51]. All participants were in the 0-14 score range, and the average ASRS score in the group was 13.15± 1.21 (n=13).

### Brain Imaging

Prior to the fMRI experiment, all participants underwent a preparatory session to verify adequate performance. During the fMRI task, participants were fitted with a response box and were directed to press the button when appropriate. Participants’ reaction times (RTs) and accuracy rates were collected.

Brain scanning was performed on a GE 3T Signa HDxt MRI scanner with an eight-channel head-coil. Functional images were acquired using a single-shot echo-planar T2*-weighted sequence. The fMRI was acquired during block-design N-back task; a working memory task known to involve the right DLPFC in correspondence to cognitive load [52], [53]. The following scanning parameters were used: TR/TE: 3000/35; flip angle 90; FOV: 20×20 cm1; matrix size: 96×96; 39 axial slices with 3 mm thickness and no gap covering the entire brain. Acquisition orientation was of the fourth ventricle plane. In addition, each functional scan was accompanied by a three-dimensional scan using T1-SPGR sequence (1×1×1 mm3).

## fMRI Analysis

Preprocessing included correction for head movement (the exclusion criterion was movements exceeding 2 mm or 2 degrees in any of the axes), realignment, normalizing the images to Talairach space and spatial smoothing (FWHM, 6 mm). The first six functional volumes, before signal stabilization, were excluded from the analysis.

Functional data were preprocessed and analyzed using BrainVoyager QX version 2.6 (Brain Innovations Maastricht, The Netherlands). The rDLPFC ROI was defined based on activity coordinates obtained from an N-back fMRI task previously performed on healthy participants in our lab [53]. We converted the statistical activation map into clusters of activation restricted by a threshold of 0.001 (bonferroni-corrected for multiple comparisons); number of voxels for the rDLPFC ROI was 10177. Whole brain statistical maps were prepared for each participant in each of the eight imaging sessions (Pre/Post for MPH/Placebo-Cognitive, MPH/Placebo-Control) using a general linear model (GLM), in which the N-back conditions were defined as predictors (0-, 2- and 3-back). In order to conduct the ROI analysis, beta values were extracted from the predefined rDLPFC area, and averaged across all voxels per participant, in each Isession.

## Statistical Analyses

As this study investigated a therapeutic effect that has not been tested before, we could not rely on prior results. Therefore, we chose to apply a within-subject design with improved statistical power (compared with the more prevalent between-subjects design), and to follow the common practice in fMRI pharmacological studies regarding sample sizes (group sizes ranging from 12 to 20 subjects; for example, see [54]–[58]; our initial sample size = 17). And indeed, our main results exhibit, in addition to significant effect sizes, substantially high statistical power, both for the brain behavior correlations (figure 3, MPH-Cog; and Figure 4a, MPH, show an observed power value of 0.94) and for the difference in improved N-Back performance between successful and unsuccessful subjects in the cognitive induction task (figure 4b; student’s t(11)= 2.48, p<0.05, Cohen’s d= 3.33).

